# Many Families of Lids for TonB-dependent Transporters in *Bacteroides*

**DOI:** 10.1101/2023.03.17.533168

**Authors:** Morgan N. Price, Adam M. Deutschbauer, Adam P. Arkin

## Abstract

Most TonB-dependent transporters in *Bacteroides* belong to the SusC family and work with an outer-membrane lipoprotein from the SusD family that functions like a lid. But the best-studied strain, *B. thetaiotaomicron* VPI-5482, also contains 18 TonB-dependent transporters that are not from the SusC family. We identified six families of putative lids for these transporters. The putative lids are lipoproteins that are encoded next to TonB-dependent transporters, but they are not homologous to SusD. AlphaFold-Multimer predicts that the putative lids bind the outer faces of the corresponding TonB-dependent receptors. Genetic data suggests that members of four of these families are required for the activity of the corresponding TonB-dependent receptors. Overall, of the 18 TonB-dependent receptors that do not belong to the SusC family, we identified alternate lids for nine.

## Introduction

In bacteria that have an outer membrane, TonB-dependent transporters (TBDTs) are transporters that can couple nutrient uptake through the outer membrane to the proton-motive force. Because TBDTs interact with molecules outside of the cell, they are also called TonB-dependent receptors. All known TBDTs contain a plug domain and a C-terminal beta-barrel domain (Noinaj et al. 2010). The beta-barrel lies within the outer membrane and forms a pore for nutrient uptake. The plug domain lies on the periplasmic side of the outer membrane and can occlude the pore. Nutrient uptake is coupled to the proton-motive force via the TonB protein, which extends across the periplasmic space and binds to the plug domain. Substrate binding to the TBDT leads to the exposure of the “TonB box” at the N-terminus of the plug domain; TonB may then drive uptake by binding the TonB box and partially unfolding the plug domain (Ratliff et al. 2021). Conformational changes to TonB are coupled to the proton-motive force via associated inner-membrane proteins ExbB and ExbD (Ratliff et al. 2021).

TBDTs are particularly abundant in *Bacteroides* and related genera. For example, while *Escherichia coli* K-12 has nine TBDTs, *B. thetaiotaomicron* VPI-5482 has 121 putative TBDTs (Pollet et al. 2021). 101 of the TBDTs in *B. thetaiotaomicron* belong to the SusC/RagA family (TIGR04056; (Haft et al. 2013)). Members of the SusC family work together with an accessory lipoprotein from the SusD family that is on the outer side of the outer membrane. In most cases, SusD proteins help to bind the substrate and are required for the receptors’ activity (Pollet et al. 2021), but there are exceptions (Phansopa et al. 2014; Huang et al. 2022; Tauzin et al. 2022). Structural studies show that SusD proteins bind to the outer face of the corresponding TBDT to enclose a large cavity, like a lid (Glenwright et al. 2017; Madej et al. 2020; Gray et al. 2021).

We noticed that many of the TBDTs in *B. thetaiotaomicron* that were *not* from the SusC family were encoded next to outer-membrane lipoproteins. And it appears that many of these lipoproteins are required for the activity of the corresponding TBDTs. So we wondered if these lipoproteins might represent additional lids, that is, proteins that bind the outer face of the TBDT and help it bind nutrients. When we used AlphaFold-Multimer (Evans et al. 2021) to predict structures for these pairs of proteins, most of these lipoproteins had high-confidence contacts with the outer face of the corresponding TBDTs, but not with other TBDTs. In total, among the 18 TBDTs of *B. thetaiotaomicron* that do not belong to the SusC family, we identified putative lids for nine of them.

## Results

### Lipoproteins associated with TBDTs in *B. thetaiotaomicron*

Of the 119 full-length TBDTs in *B. thetaiotaomicron* (Pollet et al. 2021), 101 belong to the SusC family and are encoded near a member of the SusD family. Our analysis focused on the other 18 TBDTs.

First, we searched their gene neighborhoods for lipoproteins, which we identified using SignalP 6 and PROSITE (Sigrist et al. 2013; Teufel et al. 2022). 13 of the 18 TBDTs were associated with lipoprotein(s) and were potentially cotranscribed with them (they are encoded adjacent to them and on the same strand).

We then used MicrobesOnline’s tree-browser (Dehal et al. 2010) to check if the lipoprotein was conserved near the TBDT in another family (outside of the Bacteroidaceae). 11 of the TBDTs had a conserved association with a lipoprotein (Table 1). Three of these lipoproteins were already known to be involved in nutrient uptake: BtuG2 (BT1954), XusB (BT2064), and HmuY (BT0497).

**Table 1:**
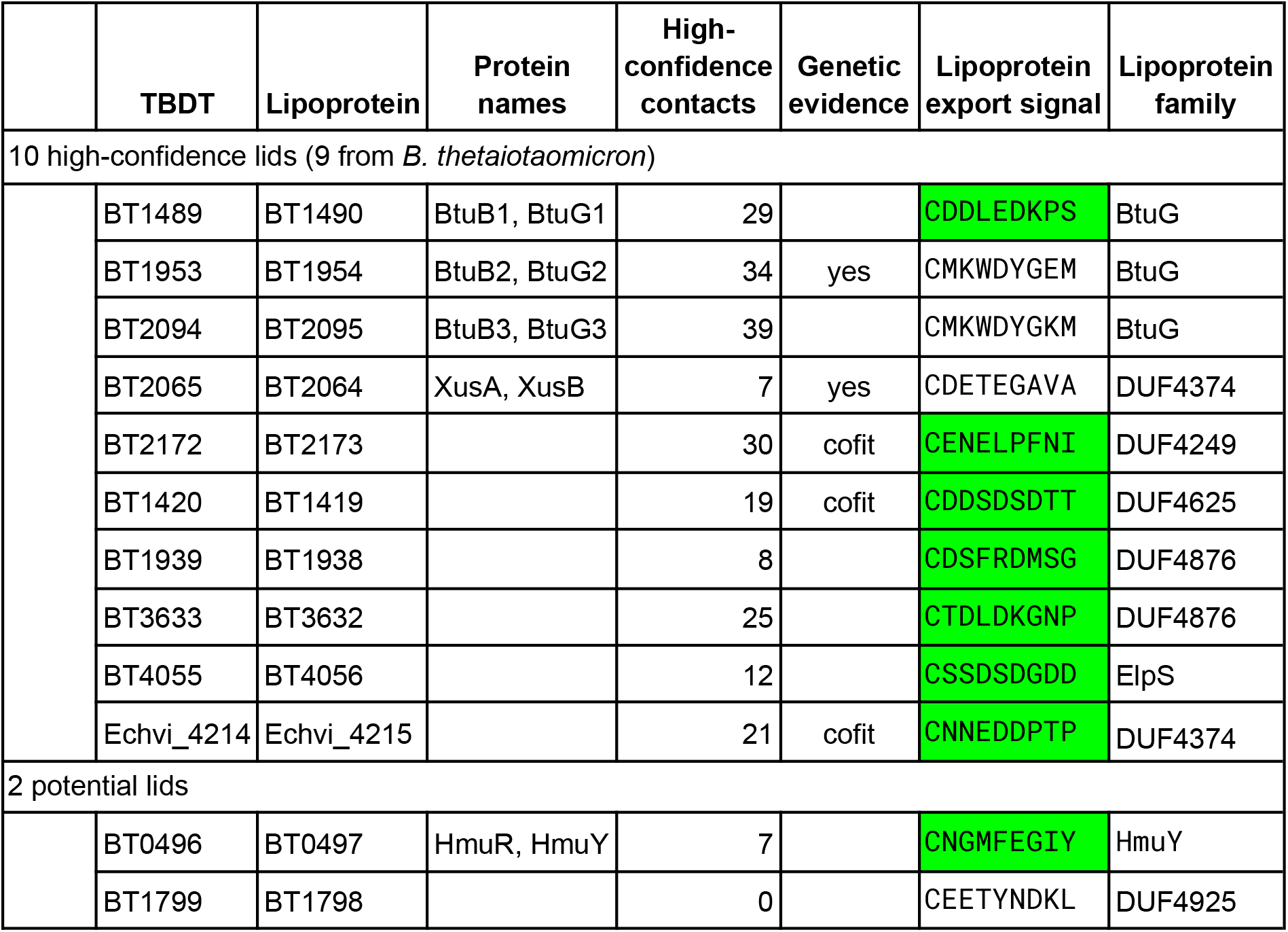
TBDTs and their associated lipoproteins. These receptors are encoded adjacent to these lipoproteins in more than one family of bacteria. High-confidence contacts are defined as pairs of residues from different chains that are predicted to be within 4Å and have a predicted aligned error (PAE) of at most 2 Å. The lipoprotein export signal is shaded if it has a bit score above zero.

BtuG2 is a surface-exposed vitamin B12-binding protein that binds the TBDT for B12 uptake, BtuB2 (Wexler et al. 2018). Furthermore, if the other B12 uptake loci are deleted, then BtuG2 is required for growth at sub-nanomolar concentrations of vitamin B12 (Wexler et al. 2018).

XusB is part of a 3-gene system for taking up enterobacterial siderophores such as enterobactin or salmochelin (Zhu et al. 2020). Mutants with any of these genes deleted are unable to grow with these siderophores as their source of iron (Zhu et al. 2020). The other components of the system are XusA (BT2065), a TBDT, and XusC (BT2063), which is related to the FoxB family of integral membrane siderophore reductases (Josts et al. 2021). (Both XusC and FoxB belong to PFam PF03929 (Finn et al. 2014).)

Finally, the amino acid sequence of BT4097 is 47% identical to that of the HmuY protein (BVU_2192) from *Phocaeicola vulgatus* (formerly *Bacteroides vulgatus*), which binds heme (Siemińska et al. 2021). A more-distant homolog from *Porphyromonas gingivalis* binds hemin (Wójtowicz et al. 2009). All of these proteins are encoded next to a TBDT known as HmuR. Strains of *P. gingivalis* with *hmuY* deleted have reduced growth with hemin or human serum as the sole source of iron; these phenotypes are similar, although perhaps less severe, to those of an *hmuR* mutant (Olczak et al. 2008).

The remaining TBDT-associated lipoproteins are two homologs of BtuB2 and six members of uncharacterized protein families (Table 1). We hypothesized that these TBDT-associated lipoproteins might function as lids and bind to the outer face of the TBDT.

### AlphaFold-Multimer analysis of potential lids

To test this hypothesis, we used ColabFold’s implementation of AlphaFold-Multimer (Evans et al. 2021; Mirdita et al. 2022) to predict the structure for each pair of proteins. We also tested a TBDT-lipoprotein pair from *Echinicola vietnamensis*, Echvi_4214 and Echvi_4215. (The rationale for studying this pair will be explained later.)

Along with the predicted structure, AlphaFold returns an estimate of the error in the pairwise distances between each pair of residues, the PAE (predicted aligned error). The PAE of the predicted interface (pairs of residues within 4 Å that are from different chains) is strongly associated with accurate predictions of protein complexes (Yin et al. 2022). So we focused on high-confidence pairs of interacting residues, which we defined as having a lowest inter-atom distance of at most 4 Å and PAE ≤ 2 both ways.

In all of the predicted complexes, the lipoprotein contacts the outer face of the TBDT. However, for BT1799 with BT1798, there are no high-confidence contacts. And for HmuR with HmuY, the only high-confidence contacts involve the portion of HmuR’s beta barrel that is expected to lie within the outer membrane (Figure 1, bottom right); none of the contacts between HmuY and the outer face of HmuR were high-confidence. The results for the other complexes support our hypothesis (Figure 1). They had 7-39 pairs of residues with high-confidence contacts (Table 1).

**Figure 1:**
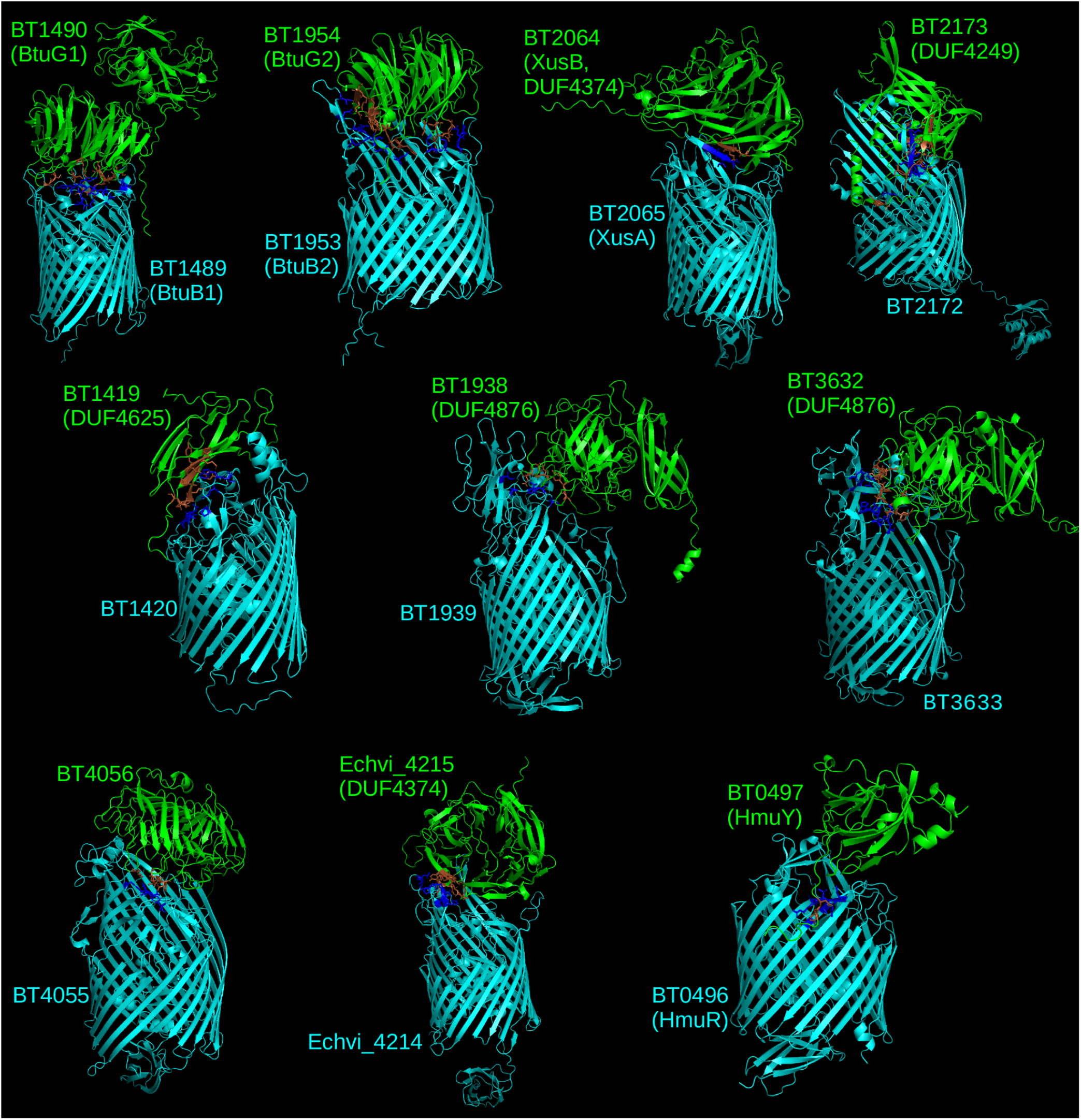
Structural predictions for lids binding to TBDTs. Residues with high-confidence contacts are shown in a darker color and with bonds between heavy atoms shown as sticks. Predictions for BtuB3/BtuG3 were similar to those for BtuB2/BtuG2 and are not shown.

As a negative control, we predicted structures for 23 random pairs of these TBDTs with other TBDT-associated lipoproteins (not the one encoded nearby). BtuB2/BtuG2 is similar to two other pairs, so we included BtuB2 and BtuG2, but not their homologs, in this test. None of the control pairs had any high-confidence contacts.

The BtuGs are predicted to form a large cavity with their TBDTs, and the cavity in the predicted structure of BtuB2/BtuG2 is large enough to contain cyanocobalamin. In most of the other structures, the lipoprotein covers most of the outer face of the TBDT, but with a channel on the side, at least 8 Å across. (This applies to XusAB, Echvi_4214-Echvi_4215, BT2172-BT2173, and BT1420-BT1419). BT4055-BT4056 has few high-confidence contacts, but BT4056 does bind on the outside, with an opening on the side of around 11 Å. In BT1939-BT1938 and BT3633-BT3632 (both lids are DUF4876), there’s a larger opening into the beta barrel (around 20 Å). Finally, the (low-confidence) structure for HmuY binding to HmuR shows two large openings on the outside.

Overall, AlphaFold-Multimer predicted that 10 of the 12 lipoproteins are likely to bind the outer face of their corresponding TBDTs. These structures are consistent with the lipoprotein helping to bind the substrate, but except for the BtuG family, the putative lids do not form a cavity. In other words, the lid is not fully closed.

### Cofitness of putative lids with their TBDTs

As discussed above, BtuG2 and XusB are already known to function with their associated TBDTs. To try and confirm a functional association between other TBDTs and their putative lids, we examined fitness data from randomly-barcoded transposon mutants of *B. thetaiotaomicron* (RB-TnSeq; (H. Liu et al. 2021)). We identified cofitness for two putative lids with their corresponding TBDTs: BT2173 (a DUF4249 lipoprotein) is cofit with the TBDT BT2172 (Figure 2A), and BT1419 (a DUF4625 lipoprotein) is cofit with the TBDT BT1420 (Figure 2B). In each case, the similarity of the genetic data for the putative lid and for the TBDT suggests that the lid is required for nutrient uptake by the TBDT.

**Figure 2:**
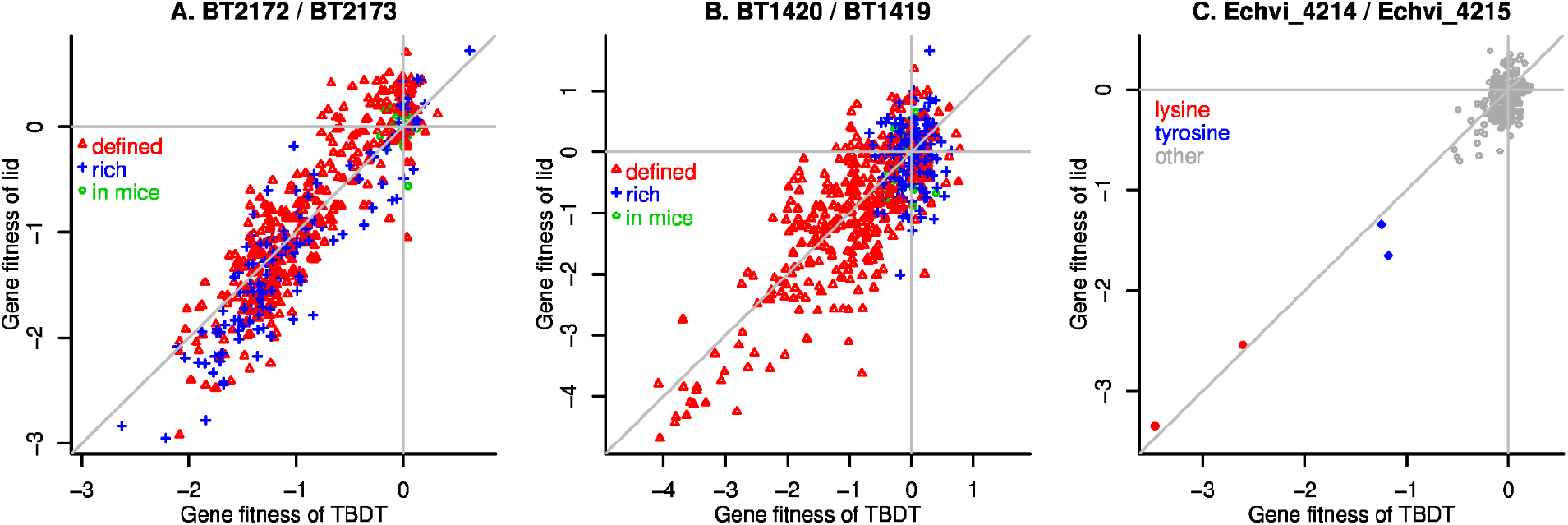
Cofitness of putative lids with their TBDTs. Panels A and B show the fitness values for the lids and TBDTs across 519 experiments for *B. thetaiotaomicron* (H. Liu et al. 2021); experiments are colored based on whether they were performed in defined media, in rich media, or in gnotobiotic mice. Panel C shows data from 202 fitness experiments for *Echinicola vietnamensis* (Price et al. 2018); experiments in defined media with lysine or tyrosine as the nitrogen source are highlighted. Except for the mouse experiments, during each experiment, a pool of barcoded transposon mutants grew from an optical density of 0.02 to saturation. In each experiment, a gene’s fitness is the log_2_ change in the relative abundance of mutants in the central 10-90% of that gene, as estimated by barcode sequencing (Wetmore et al. 2015). The lines show *x* = 0, *y* =0, and *x* = *y*.

Also, while doing this analysis, we noticed that a distant homolog of the XusA-XusB pair, Echvi_4214 and Echvi_4215 from *Echinicola vietnamensis*, have similar fitness patterns (Figure 2C; data of (Price et al. 2018)). Both genes are important for the utilization of lysine or tyrosine as the nitrogen source. Echvi_4214 is 30% identical to XusA, while Echvi_4215 is distantly related to XusB. XusB belongs to DUF4374 (PFam PF14298); Echvi_4215 is not assigned to DUF4374 by PFam’s hidden markov model, but some BLAST-level homologs are, and Echvi_4215 is assigned to this family by a deep learning approach (PFam-N, (Bileschi et al. 2022)). AlphaFold predicts fairly-similar structures for XusB and Echvi_4215 (root mean square distance of 3.02 and template modeling score of 0.48 over 54-57% of the proteins, despite 13% sequence identity; structures were taken from the AlphaFold protein structure database (Varadi et al. 2022)). Echvi_4214 and Echvi_4215 are encoded next to a distant homolog of XusC, Echvi_4216 (28% identity), but mutants of Echvi_4216 did not have any phenotypes (all 202 fitness values were within ±0.5). Overall, it appears that the putative lid Echvi_4215 is required for nutrient uptake by Echvi_4214.

If we combine the cofitness results with the prior studies of BtuB2/BtuG2 and XusA/XusB, and we extend the functional relationship of BtuB2/BtuG2 to the two paralogous pairs, then we have genetic evidence of a functional relationship for six of the nine putative TBDT/lid pairs from *B. thetaiotaomicron*. None of the genes in the other three pairs had any clear phenotypes. (All 6 * 519 fitness values were within ±1 except for BT3632, which has a value of −1.3 in one replicate of dimetridazole stress at 0.025 mM, but has values of 0.0 and −0.3 in the other two replicates of this experiment.)

### Conserved proximity of TBDTs and putative lids

For the three pairs without any genetic evidence for a functional relationship, we used *fast.genomics* (http://fast.genomics.lbl.gov) to check how conserved their proximity was. As shown in Figure 3, all three pairs are found near each other and on the same strand in representative genomes from dozens of genera. (Homologs of BT1939 and BT1938 were found nearby in 81 genera; BT3633/BT3632 were nearby in 59 genera; and BT4055/BT4056 were nearby in 621 genera.) Homologs with above 20% of the maximum bit score were almost always close to each other (Figure 3). Such conserved proximity supports our hypothesis that they interact physically (Dandekar et al. 1998).

**Figure 3:**
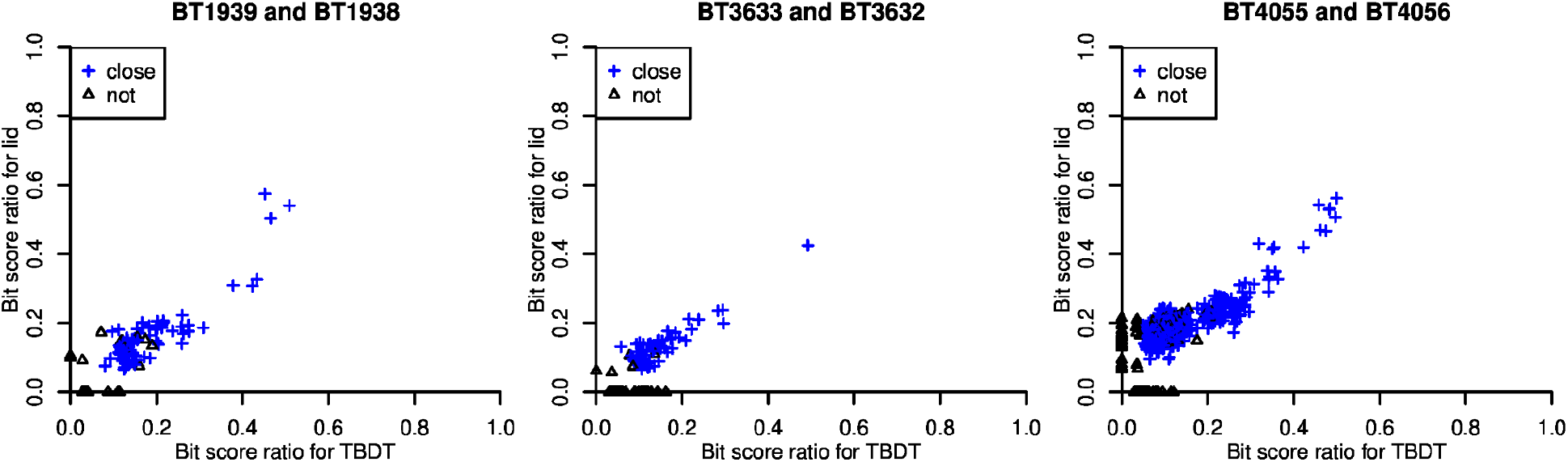
Conserved proximity for three TBDTs and their putative lids. Each panel shows a different pair, and each point shows a genome that contains a homolog (E < 0.001) of the TBDT and/or the lid. The *x* axis shows the bit score ratio (relative to the maximum possible) for the TBDT, or zero if no homolog was found. Similarly, the *y* axis shows the bit score ratio for the lid. Points are colored blue if the highest-scoring homologs to the TBDT and lid are separated by 5 kb or less and are on the same strand. *Fast.genomics* includes one representative genome each for 4,788 genera of bacteria and archaea, so every genome is from a different genus (as classified by GTDB (Parks et al. 2018)).

### Localization of the putative lids

For the lipoproteins to function as lids, they need to reach the outer face of the outer membrane. To test if the putative lids are localized to the outer membrane, we used two strategies: we searched for experimental data as to the localization of the lids or their homologs, and we analyzed the sequences of the lids for lipoprotein export signals (Lauber et al. 2016; Valguarnera et al. 2018).

As far as we know, the only putative lid that has been shown to be surface exposed is BtuG2; specifically, BtuG2 is degraded upon exposure of intact cells to a protease (Wexler et al. 2018). BtuG2’s paralogs, BtuG1 and BtuG3 (BT1490 and BT2095), are likely to be on the outer side of the outer membrane as well. Three other potential lids have homologs that are reported to be on the surface. First, BT1938 is 30% identical to BF9343_2981 from *B. fragilis*, which was sensitive to the treatment of intact cells with proteinase K and was labeled by a biotinylation reagent (Wilson et al. 2015). Second, BT4056 is 40% identical to Fjoh_0546 from *Flavobacterium johnsoniae*, which was enriched in the secreted protein fraction (“exoproteome”) after phosphate depletion (Lidbury et al. 2021). This is consistent with an outer-facing localization of Fjoh_0546 because an outer-facing lipoprotein is tethered to the membrane only by its lipid anchor, while an inner-facing lipoprotein would be more likely to be retained within the cell. BT4056 is also distantly related to ElpS from *Caulobacter crescentus*, which is secreted; it is thought that ElpS is an outer-facing lipoprotein before proteolytic cleavage (Le Blastier et al. 2010). Third, HmuY (BT0497) is 48% identical to BF9343_2622 from *B. fragilis*, which is reported to be secreted (Wilson et al. 2015). Furthermore, a distant homolog of HmuY, RhuA from *Riemerella anatipestifer*, was shown to be present on the cell surface by immunolabeling (M. Liu et al. 2021).

Another way to predict the localization of the putative lids is from the sequences just after the N-terminal cysteine (which is formed by cleaving off the lipoprotein attachment signal). In Bacteroidota, these sequences are known as lipoprotein export signals and they drive the export of lipoproteins to the outer face of the outer membrane (Lauber et al. 2016; Valguarnera et al. 2018). To identify sequences that are likely to function as a lipoprotein export signal, we used a position weight matrix approach with two different training sets. Our first training set was 88 SusD proteins from *B. thetaiotaomicron*, as these proteins are expected to be outer facing. (This is the same strategy as used previously to identify the lipoprotein export signal in *Flavobacterium johnsoniae* (Lauber et al. 2016).) Our second training set was 91 lipoproteins that are found in outer membrane vesicles from *B. thetaiotaomicron* (Valguarnera et al. 2018); the rationale for this training set is that these lipoproteins are enriched in export signals (Valguarnera et al. 2018). This gave us two position weight matrices (Figure 4A and 3B). Both weight matrices gave a strong discrimination between their training sets and random subsequences of lipoproteins from *B. thetaiotaomicron*, and bit scores above zero are far more likely for sequences from the training set (Figure 4C).

**Figure 4:**
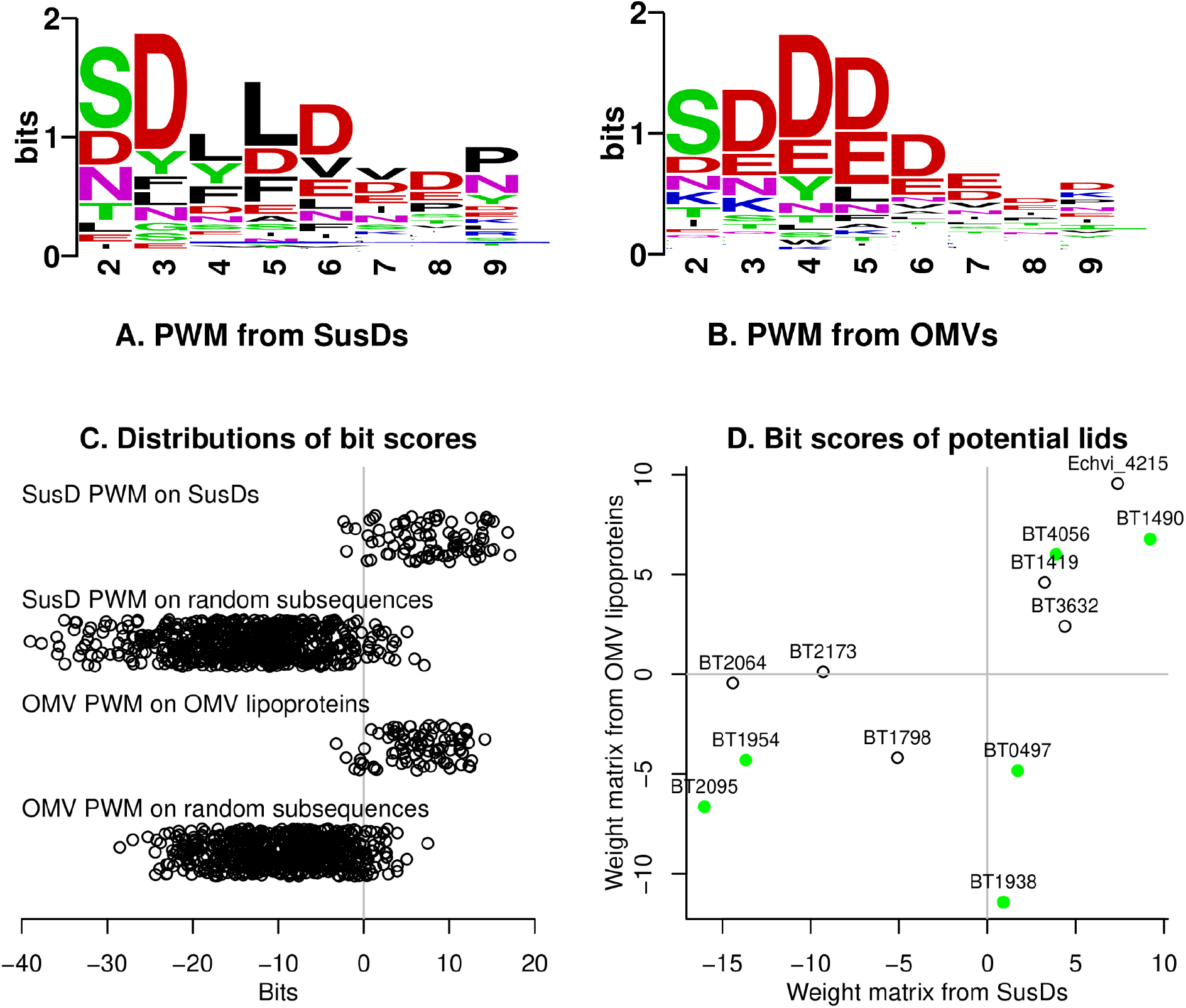
Analysis of lipoprotein export signals. (A) Sequence logo for the position weight matrix derived from SusD proteins (drawn with weblogo, (Crooks et al. 2004)). The weight matrix begins immediately after the N-terminal cysteine. (B) Sequence logo derived from outer membrane vesicle lipoproteins. (C) Distribution of bit scores for training sequences and random 8-mers from *B. thetaiotaomicron* lipoproteins. (D) Bit scores for the export signals in the potential lids. Proteins whose homologs are known to be exported are highlighted (green filled dots).

When we applied these weight matrices, to the putative lids, we found that most of the lids had a plausible match (bits > 0) for one of the two weight matrices (Figure 4D). Export signals with positive scores are also highlighted with a green background in Table 1; the invariant N-terminal cysteine is shown there but is not included in the scoring. Two of the lids without positive scores were BT1954 (BtuG2), which is known to be surface exposed (Wexler et al. 2018), and its paralog BT2095 (BtuG3), which is also probably exported. The two proteins have nearly identical export signals (Table 1) and we wonder if they are exported by another mechanism. The other exceptions were BT2064 (XusB) and BT1798. Some close homologs of BT2064 have high-scoring export signals, and its OMV score was near zero, so we do expect that it is outer-facing. (For instance, BDI_3402 from *Parabacteroides distasonis* is 61% identical and has an export signal of CSDDDPVNP, with bit scores of +8.7 or +12.2.) BT1798 did not have any high-confidence contacts with the associated TBDT, and it might be in the periplasm.

Overall, we identified an exported homolog or a plausible lipoprotein export signal for all of lids with high-confidence contacts with their TBDTs, with the potential exception of XusB.

### Lid-associated TBDTs are no longer than TBDTs without known partners

TBDTs from the SusC family tend to be longer (have more amino acids) than other TBDTs, presumably because longer loops on the outside of the beta barrel interact with their partner proteins (Pollet et al. 2021). So we wondered if our putative lid-associated TBDTs had long beta-barrel domains. To estimate the size of these domains, we used the number of amino acids C-terminal to the plug domain (PFam PF07715). As shown in Figure 5, SusC family TBDTs from *B. thetaiotaomicron* have substantially longer beta-barrel domains. But lid-associated TBDTs are about as long as TBDTs that do not have an apparent lipoprotein partner (medians of 569 and 567 amino acids, respectively). The characterized TBDTs from *E. coli* have similar lengths for their beta barrel domains (468 to 592 amino acids).

**Figure 5:**
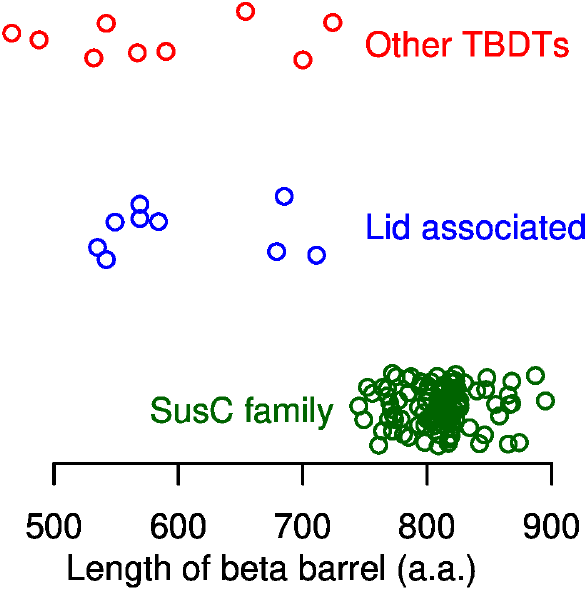
Length of the beta barrel region in 119 TBDTs from *B. thetaiotaomicron*. The beta barrel region was defined as from the end of the plug domain to the end of the protein.

## Discussion

### TBDTs with alternate lids have smaller substrates than SusC-like transporters

Most SusC-like transporters act on oligosaccharides or peptides. This makes sense given that SusCD complexes have large cavities that hold multiple 2.5 kD oligosaccharides or peptides of up to 13 amino acids or around 1.4 kD (Glenwright et al. 2017; Madej et al. 2020; Gray et al. 2021). In contrast, all of the known substrates for TBDTs with alternate lids are smaller. BtuB2 transports cyanocobalamin (1.4 kD), and BtuB1 and BtuB3 are also expected to transport cobamides. XusA transports enterochelin and salmochelin (0.6-0.7 kD). Echvi_4214 appears to transport lysine (although the gene cluster also contains a potential siderophore reductase, so we are not sure if lysine is the physiological substrate). Acting on smaller substrates may explain why these TBDTs lack the long extracellular loops of SusC proteins.

### What is the role of the lids?

Most of the putative lids belong to families about which little is known: BT1419 belongs to DUF4625; BT1938 and BT3632 belong to DUF4876; XusB and Echvi_4215 belong to DUF4374; and BT4056 does not belong to a PFam, but InterPro places it in the pectin lyase superfamily (Finn et al. 2017). The most obvious role for the lids would be to help bind the substrate, and there is evidence of substrate binding for BtuG2 and HmuY (although AlphaFold did not find high-confidence contacts for HmuY with HmuR). More speculatively, the lids could signal substrate binding to the TBDT and facilitate the exposure of the TonB box, as occurs for SusD proteins (White et al. 2022).

### Why are lids associated with Bacteroidota?

For most of the pairs of TBDTs with their putative lids, virtually all of the homologs with conserved proximity are in Bacteroidota. The exception is BT4055/BT4056, which has homologs in proximity in 621 genera, including 394 genera of Bacteroidota and 171 genera of Proteobacteria. For example, ElpS from *Caulobacter crescentus* (an α-Proteobacterium) is an outer-facing lipoprotein (Le Blastier et al. 2010) that is homologous to BT4056, and ElpS is encoded adjacent to TonB-dependent receptor CCNA_00170. SusC and SusD are also found primarily in Bacteroidota (352/357 genera with homologs near each other are from Bacteroidota). Our best guess is that most other diderm bacteria lack the machinery to export these lipoproteins to the outside of the cell, but this doesn’t seem consistent with the broad taxonomic distribution of ElpS.

## Materials and Methods

### TonB-dependent transporters and associated lipoproteins

121 TBDTs in *B. thetaiotaomicron* VPI-5482 were identified previously (Pollet et al. 2021). After excluding two TBDTs with putative frameshifts (BT3493/3494 and BT3750/3751), we had 119 full-length TBDTs to analyze. We found three additional proteins in the genome that contained the plug domain (BT1139, BT4200, and BT4324 contain PFam PF07715), but their AlphaFold predictions did not show a beta-barrel domain, so we did not consider them further. SusC-type TonB-dependent receptors were identified by using TIGRFam TIGR04056 (Haft et al. 2013) with the trusted cutoff.

We identified potential lipoproteins that were encoded adjacent to and on the same strand as non-SusC-type TonB-dependent receptors by using SignalP 6 and PROSITE (Sigrist et al. 2013; Teufel et al. 2022). These tools search for a “lipobox” motif C-terminal to the signal peptide. Although the annotated protein sequence for BT1938 has no signal peptide or lipobox, we found homologs of BT1938 that are longer and are annotated as lipoproteins. We then noticed that BT1938 has an upstream ATG; translation from this start codon adds an additional N-terminal sequence that contains a signal peptide and a lipobox (MKKYLIYLFTLASTLLIGCDSFRD).

Besides the pairs listed in Table 1, the TBDT BT0150 is encoded next to lipoprotein BT0149. However, BT0149 is probably in the periplasm: it is 40% identical to LoiP from *E. coli*, a protease that is thought to be located on the inner face of the outer membrane (Lütticke et al. 2012). Furthermore, the bit scores of BT0149’s export signal were strongly negative (−21.5 and −15.8). Also, BT0149’s potential function as a protease is not consistent with acting as a lid. We did consider this pair any further.

### ColabFold

We ran ColabFold (Mirdita et al. 2022) with default settings via google CoLab Pro+. These settings include using both paired and unpaired alignments, 5 models per pair, and no refinement with amber. For the lipoproteins, we removed the signal peptide and the lipobox so that the sequence began with the mature cysteine. For BT1938, we included the additional sequence that is upstream of the annotated ATG but after the lipobox (CDSFRD).

### Analysis of conserved proximity with *fast.genomics*

*Fast.genomics* will be described in more detail elsewhere. It contains one high-quality representative genome for each of 4,788 genera, as identified using GTDB release 207 (Parks et al. 2018) and GUNC (Orakov et al. 2021). Homologs were identified using mmseqs2 (Steinegger and Söding 2017), with a sensitivity setting of 7 for proteins with ≤150 amino acids or 6 otherwise.

### Analysis of lipoprotein export signals

To build position weight matrices of lipoprotein export signals, we used outer membrane vesicle proteins from *B. thetaiotaomicron* (Valguarnera et al. 2018) or the SusD proteins from *B. thetaiotaomicron* (listed by (Pollet et al. 2021)). For the outer membrane vesicle proteins, we used the export signal sequences previously reported. For the SusD proteins, we used SignalP 6.0 (Teufel et al. 2022) to identify the processed cysteine. SignalP identified the lipobox for only 88 of 98 SusD proteins, so we used those 88 sequences. To describe the background distribution, we used the frequencies of the amino acids across all predicted proteins in *B. thetaiotaomicron*. For the weight matrix, we used eight positions after the N-terminal cysteine, because subsequent positions showed little difference from the background distribution.

The bit score or weight for each amino acid at each position is log_2_(frequency in training set / background frequency). To compute the frequencies, we used a pseudocount of 1 (we added 1 to the count of each amino acid at each position). The bit score for a potential lipoprotein export sequence is the sum of the bit scores for that sequence’s amino acid at each position.

## Acknowledgments

This material by ENIGMA-Ecosystems and Networks Integrated with Genes and Molecular Assemblies (http://enigma.lbl.gov), a Science Focus Area Program at Lawrence Berkeley National Laboratory is based upon work supported by the U.S. Department of Energy, Office of Science, Office of Biological & Environmental Research under contract number DE-AC02-05CH11231

